# Muscle mass and denervation explain variability in maximal power and rapid force across the adult female lifespan

**DOI:** 10.64898/2026.07.13.738367

**Authors:** Steven J. O’Bryan, Annabel Critchlow, Andrew Garnham, Christopher S. Fry, Danielle Hiam, Séverine Lamon

**Author notes:** denotes equal contribution (Danielle Hiam, Séverine Lamon). **Corresponding author:** Steven J. O’Bryan, 0399195806.

## Abstract

**Background:** Dynamic power declines earlier across the lifespan and shows a more pronounced and complex pattern than isometric strength, particularly in ageing females. However, the functional, skeletal muscle and molecular mechanisms underpinning power loss across the female lifespan remain to be collectively examined.

**Methods:** Eighty-six females aged 18-80 years and stratified per decade of age completed a series of maximal voluntary knee extensions to construct torque-velocity and power-velocity relationships of the quadriceps. Data points corresponding to >95% maximal power were selected for the evaluation of rate of torque development (RTD) and quadriceps surface electromyography (EMG). Outcomes were quantified within discrete 50ms time bins from torque onset to +200ms and included absolute RTD, RTD normalised to peak force, and EMG amplitude and rate of rise normalised to the maximal compound action potential. Quadriceps morphology was assessed via computed tomography, and a *vastus lateralis* muscle biopsy was collected to assess markers of denervation and expression of genes associated with the neuromuscular junction and calcium-handling transcriptome.

**Results:** Ageing led to linear reductions in maximal power (-1.39 ± 0.01% p/year), torque (-0.98 ± 0.13% p/year) and velocity (-0.38 ± 0.01% p/year) (all p < 0.05). Quadriceps skeletal muscle CSA attenuated power loss by ∼40% (p < 0.001), largely through reduction of the decline in torque (∼50%), with no effect on the decline in velocity. During early time bins, older females generated higher relative RTD accompanied by higher EMG amplitude, whereas during later time bins, older females generated less absolute and relative RTD accompanied by lower EMG amplitude and rate of rise (all p < 0.05). Ageing increased neural cell adhesion molecule (NCAM) positive fibres and fibrosis (both p < 0.05). The presence of NCAM□ fibres was associated with attenuation of the age-related decline in maximal power (∼15%), torque (∼35%) and velocity (∼60%), suggesting that NCAM□ fibre prevalence may partially explain the observed age associations. Within the neuromuscular junction transcriptome, ageing reduced acetylcholinesterase and increased laminin alpha-2 and muscle-specific kinase (all FDR < 0.05), whereas lesser changes were observed within the calcium-handling transcriptome.

**Conclusions:** Skeletal muscle CSA explains ∼40% of the age-related decline in quadriceps dynamic maximal power across the female lifespan, whereas a neurodegenerative profile mainly evidenced by age-related changes in voluntary neural drive, denervation and markers of neuromuscular junction instability further contribute to the decline.

## 1. INTRODUCTION

Epidemiological studies indicate that the severity and prevalence of sarcopenia are higher in ageing males when defined using criteria based on handgrip strength and appendicular skeletal muscle mass (e.g. EWGSOP2), whereas definitions that also consider gait speed (e.g. IWGS) show similar or higher rates and severity in ageing females [1]. Despite the global use of hand grip strength as a general health biomarker and its ease in measurement, its direct relevance to lower-body physical function and inclusion as the sole measure of muscle strength used in sarcopenia diagnosis has been questioned [2]. Indeed, quadriceps strength is more closely related to lower-body physical function and mobility throughout the lifespan compared to handgrip strength [3]. Notably, females experience a greater decline in quadriceps strength from 40-50 years of age compared to males [4]. Further, primary outcomes derived from maximal dynamic muscle actions including power production and velocity of movement have negative and independent associations with physical disability [5, 6], exhibit higher hazard ratios for mortality [7] and are more sensitive to age-related changes in neuromuscular function compared to isometric strength tests [3, 8].

Recently we demonstrated accelerated reductions in quadriceps isometric maximal voluntary and evoked torques and estimated one-repetition maximum (e1RM) during the menopausal transition, with quadriceps skeletal muscle cross-sectional area being relatively more important for isometric torque production compared to the e1RM [9]. These findings support different mechanisms and trajectory in the age-related decline in isometric torques versus dynamic strength and power [10], and others have reported more rapid and severe decreases in crude power compared to isometric and dynamic torques [3, 4]. Age effects on maximal dynamic power are most rigorously examined by quantifying changes in the hyperbolic/linear/quasi-linear torque-velocity relationship and the parabolic power-velocity relationship from whole muscle/limb maximal voluntary contractions performed under various isotonic loads [11], with such studies attributing quadriceps peak power loss in middle-aged and older females to decreases in torque capacity followed by contraction velocity [6]. Our recent findings suggest that peripheral mechanisms, evidenced by decreases in high and low frequency paired pulse responses, primarily contribute to reduced isometric strength across the female lifespan, and that *rectus femoris* quadriceps muscle may be more susceptible to age-related denervation than the *vastu*s quadriceps muscles [9]. However, how these changes influence dynamic torque-velocity and power-velocity outcomes and neural drive to quadriceps at different stages of the female lifespan remain unknown.

Rate of torque development (RTD) is a key determinant of outcomes derived from the quadriceps torque-velocity relationship [12] and plays a key role in correcting balance perturbations, minimizing falls risk and associated hospitalizations, and increasing gait speed in older adults [13]. Further, voluntary RTD during earlier time bins from torque onset up to ∼100ms are mostly sensitive to motor unit recruitment, whereas later time bins between 100-250ms post torque onset are mostly sensitive to intrinsic contractile properties (e.g. Ca^2+^ kinetics/sensitivity and fibre composition/properties), skeletal muscle morphology (e.g. including cross-sectional area and intramuscular adiposity) and musculotendinous stiffness [14–16]. Although absolute RTD decreases with ageing and likely contributes to power loss, normalisation to peak force removes the age effect and has been attributed to differences in skeletal muscle mass [17, 18]. However, the relationship between skeletal muscle CSA and voluntary strength is not perfectly linear, with skeletal muscle CSA explaining ∼65% of the variance in isometric force across the female lifespan [19]. Hence, comparisons of statistical models adjusted for skeletal muscle mass rather than solely normalising to peak force provides a better approach for understanding its contribution to functional decline with ageing.

At the myocellular level, we recently reported ageing females showed no changes in *vastus lateralis* type IIa fibre CSA or proportion, a decrease in type I fibre CSA and an increase in hybrid type I/IIa proportion [20]. An increase in hybrid fibres may represent an age-related decrease in alpha motoneuron size/number or neuromuscular junction instability leading to collateral axonal sprouting to adjacent fibres [21]. These changes can occur prior to any changes in fibre size and may explain the disproportionate age-related decline in muscle strength/power compared to mass [3]. Potential destabilisation of the neuromuscular junction and denervation of skeletal muscle fibres in human ageing can be reflected by the gene or protein expression of several molecular markers including neural cell adhesion molecule (NCAM), acetylcholine receptors (AChR α, β, δ, γ) and muscle-specific kinase (MuSK) [22], many of which are upregulated in response to neuromuscular junction instability and denervation in older skeletal muscle [23]. Beyond changes in fibre morphology/proportion and denervation, other mechanisms likely to contribute to reduced power and rapid force in female ageing includes changes in intrinsic muscle function [9, 10] reflected by changes in the calcium handling transcriptome [24] and increased intermyofibrillar fibrosis and adiposity [20, 21].

The aims of this study were to map the trajectory and severity of age-related changes in force-velocity, power-velocity and rapid force characteristics of the quadriceps across each decade of the adult female lifespan, and to explore potential relationships with changes in skeletal muscle mass, voluntary neural drive, denervation and the neuromuscular junction and intrinsic calcium handling transcriptome. The findings from this study may provide direct mechanistic evidence and focus of intervention for improving functional capacity in ageing females and reducing sarcopenic incidence.

## 2. METHOD

### 2.1 Ethical approval and participants

This study was approved by Deakin University’s Human Research Ethics Committee (DUHREC 2021-307) and conducted in accordance with the standards set by the Declaration of Helsinki, except for registration in a database. Participant recruitment including inclusion/exclusion criteria has already been presented [9, 20]. Eighty-six females evenly represented over each decade provided written informed consent and completed the study protocols.

### 2.2 Experimental protocol and procedures

The experimental protocols have previously been described in detail [9, 20, 25] and can also be found in the Supporting Information.

### 2.3 Data analysis

#### 2.3.1 Torque-velocity and Power-velocity relationships

Several modelling methods were applied to both average and peak data with that returning the highest adjusted R² and lowest standard error of the estimate (SEE) chosen. For torque-velocity (T-V) this included comparing linear regression and Hill’s hyperbolic least squares method and for power-velocity (P-V) this included comparing Hill’s re-organized equation and quadratic and cubic polynomial regressions with intercept set at zero [11, 26]. On this basis, peak T-V were best modelled using Hill’s hyperbolic least square method for all participants (Figure 1A) and peak P-V was best fit via cubic polynomial regression (Figure 1B). Both efforts completed at each load/velocity were included in modelling procedures unless they deviated by >5% from each other (in which case only the highest value was included) and the following variables were extracted: maximal torque (T_0_) at the y-axis intercept of the T-V relationship, 2) maximal velocity (V_0_) at the x-axis intercept of the T-V relationship, 3) maximal power (P_MAX_) at the apex of the P-V relationship, and 4) optimal velocity (V_opt_) at corresponding P_MAX_.

**Figure 1.**
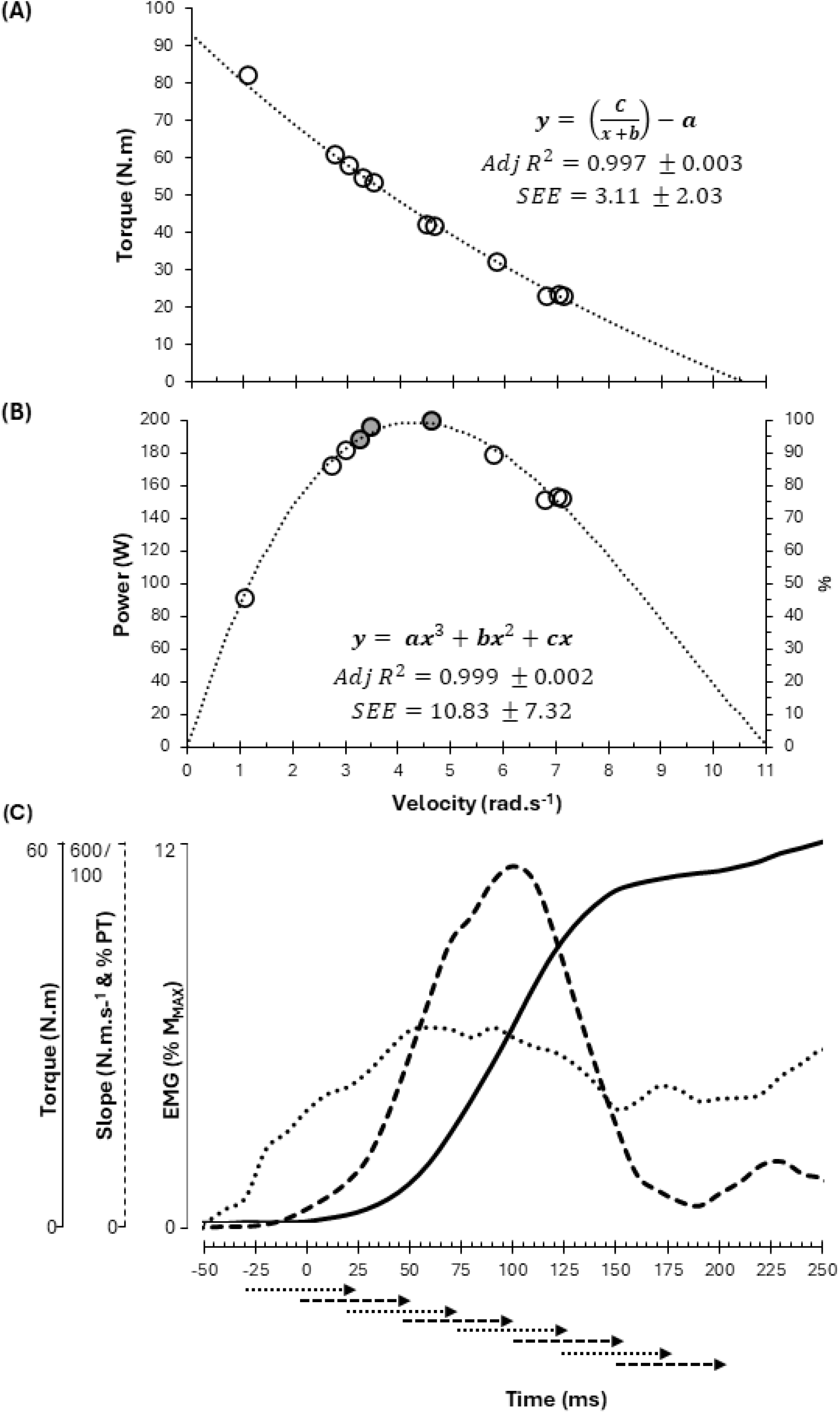
**(A)** Peak knee extension torque and **(B)** power plotted against peak velocity for maximal isoinertial (0 up to 65% MVC) and isokinetic (1.047 rad.s^-1^) contractions for a representative participant. T0 and V0 were extracted as the y and x intercepts of the T-V relationship, and P_MAX_ and V_opt_ were extracted as the apex of the curve and corresponding velocity from the P-V relationship. The modelling equations and overall group mean ± SD for the adjusted R^2^ and standard error of the estimate (SEE) are shown. **(C)** Power values =>95% P_MAX_ (grey fill) were selected for further analysis including rate of torque development (RTD) and quadriceps neural drive (4 ± 2 data points per participant). Torque onset was set at ∼5% peak slope (0ms) and EMG onset as ∼0.02mV increase above baseline for > 5ms (both visually inspected) with the time difference between onsets representing the electromechanical delay (EMD). From the torque slope and torque-time curve, maximum RTD, average RTD and total impulse during 50ms time bins from torque onset to plus 200ms were calculated. Similarly, for each quadriceps muscle RMS average amplitude, maximum amplitude and rate of EMG rise were calculated across 0-50ms and 50ms time intervals from EMG onset to plus 200ms (+100ms for RER) and normalised to maximal M-wave amplitude.

#### 2.3.2 Rate of torque development and electromyography

Data points from the P-V relationship which fell within ≥95% of P_MAX_ were chosen for rate of torque development analysis (4 ± 2 data points per participant) (Figure 1C). The slope of the torque-time curve (10ms constant) was used to determine maximal voluntary RTD (peak slope value) and average RTD in absolute (N·m.s^-1^) and relative (normalized % peak voluntary torque) terms during time bins including 0-50ms, 50-100ms, 100-150ms and 150-200ms from torque onset [17, 18, 27]. Torque onset was identified by an automatic cursor set at rising threshold of 5% of the peak slope, viewed in conjunction with the original unprocessed torque signal in high resolution (∼5 N·m and 25ms) and then manually adjusted if necessary [27].

To determine muscle activation onsets, the filtered EMG signal was rectified and smoothed (2.5ms root mean squared) and a cursor was automatically set at a rising threshold of ∼0.02mV above baseline for > 5ms, visually inspected in high resolution alongside the torque signal and torque onset cursor, and manually adjusted if necessary. From EMG onset to offset knee angle, a 50ms root mean square (RMS) was applied to EMG signals then normalised to the peak-to-peak amplitude of the maximal M-wave elicited at 70° knee flexion (i.e., mid-range) immediately following a pair of contractions performed at a given isotonic load (%M_MAX_). The average, peak and rate of EMG rise (EMG slope (RER)) for each muscle were calculated for the same time bins as RTD (except for RER which was calculated from up to 100ms only) and individually offset for the electromechanical delay. To represent muscle activation during the same time bins as RTD, time bin EMG outcomes were normalized to M_MAX_ elicited at 110° knee flexion to ensure comparable spatial representation of sampled motor units.

#### 2.3.3 Immunohistochemical staining

*Vastus lateralis* muscle samples from a subset of 33 randomly selected participants (n = 6 for 18-29, 30-39, 40-49, 60-69; n = 5 for 50-59; n = 4 for 70-80) were cut into 10µm sections using a cryostat and stored at -80°C. To quantify denervation, the muscle sections were fixed in ice cold acetone for 3 minutes, washed in phosphate-buffered saline (PBS), incubated in 2.5% normal horse serum for one hour, then incubated overnight with α-NCAM (neural cell adhesion molecule, 1:50, BD Biosciences, Franklin Lakes, NJ, USA) and α-laminin (1:100, Merck, Burlington, MA, USA). Following a wash in PBS, the sections were incubated for 75 minutes with goat α-mouse Alexa Fluor 555 (1:250, Thermo Fisher Scientific, Waltham, MA, USA) and goat α-rabbit Alexa Fluor 488 (1:250, Thermo Fisher Scientific), then washed again. The sections were incubated for 10 minutes with DAPI (1:10,000, Invitrogen, Waltham, MA, USA), washed in PBS, and mounted with Vectashield mounting media (Vector Laboratories, Newark, CA, USA). Finally, the sections were imaged at 10 x magnification using the tiles and stitching functions on an AxioImager M2 upright microscope (Zeiss Zen 3.1, Oberkochen, Germany; Figure 1). Fibres were deemed dennervated (NCAM^+^) if NCAM was strongly expressed throughout the entire fibre. NCAM^+^ fibres were expressed as a percentage of total fibre number.

#### 2.3.4 RNA sequencing

A subset of 66 participants underwent muscle RNA sequencing, as previously published [20, 28]. RNA was extracted from 10mg of snap frozen *vastus lateralis* tissue, using the AllPrep DNA/RNA/miRNA Universal kit (Qiagen, Hilden, Germany). cDNA libraries were prepared using the Ribo-Zero Gold kit and the Illumina TruSeq Stranded Total RNA protocol (Illumina, San Diego, CA). RNAseq libraries were prepared using the Illumina TruSeq Stranded Total RNA with Ribo-Zero Gold protocol and sequenced with 150-bp paired-end reads on the Illumina Novaseq6000 (Macrogen Oceania Platform). Reads underwent quality check with FastQC (v0.11.9; Babraham Institute, Cambridge, UK) and Kallisto (v0.48.0) was used to map reads to the human reference genome (*HomoSapien GRCh38)* and to generate transcript counts. One sample was excluded from analysis as it did not pass quality control due to poor alignment. The counts matrices from the three batches of samples were merged into one matrix. The complete set of results can be found on the Gene Expression Omnibus (GEO) platform under accession number GSE303107. Targeted analysis of a subset of genes representing the regulation of NMJ signalling, NJM structure and function, ECM regulation, fibrosis and Ca^+2^ handling was undertaken for the purpose of this study. All genes investigated are represented in Supporting Information.

### 2.4 Statistics

All analyses were conducted in R (version 4.4.3 (2025-02-28)), with statistical significance set at *p* < 0.05. Additional statistical information is presented in Supporting Information.

To examine associations between age and each outcome, linear regression models were initially fitted (*Outcome* ∼ *Age* + *MVPA* + *Protein*). Generalised additive models (GAMs) were then fitted to assess non-linear age associations using thin plate regression splines with smoothing parameters estimated using restricted maximum likelihood (REML) [29]. Time-series effects were examined using mixed-effects models with the lme function. The fixed-effects structure included TimeBin, Age, and their interaction and covariates (MVPA and protein intake). Secondary models were additionally adjusted for quadriceps CSA to determine whether associations were independent of muscle size. Random intercepts and random slope models were compared using AIC. The TimeBin x Age interaction was assessed using Type III ANOVA for significance. Where significant, post hoc analyses were conducted using estimated marginal trends (*emtrends)*, representing the association between age and the outcome at each time bin, with interpretation based on slope estimates and 95% confidence intervals. Pairwise comparisons of slopes between time bins were then performed using *pairs*, with Tukey adjustment for multiple comparisons, to determine whether the strength of the age–outcome association differed between phases.

Differential expression analyses were performed transcriptome-wide however, interpretation and downstream visualisation focused on genes associated with neuromuscular junction and calcium handling pathways. Differential expression analysis was performed using the DESeq2 package (v1.46.0) [30]. To identify age-associated changes in gene expression, negative binomial generalised linear models were fitted using the design formula:∼ batch age where age was treated as a continuous variable and sequencing batch was included as a covariate. Differentially expressed genes were identified using Wald tests, with statistical significance determined using Benjamini–Hochberg adjusted p-values (FDR < 0.05).

The code for this analysis can be found at https://github.com/DaniHiam/FAMe-Neural-drive-and-the-neuromuscular-transcriptome-.

## 3. RESULTS

### 3.1 Participant characteristics

Participant characteristics are reported in table 1.

**Table 1.** Participant characteristics.

| Variable | Overall | 18-29<br>N = 19 | 30-39<br>N = 13 | 40-49<br>N = 12 | 50-59<br>N = 14 | 60-69<br>N = 15 | 70-80<br>N = 13 |
| --- | --- | --- | --- | --- | --- | --- | --- |
| **Participant characteristics** |  |  |  |  |  |  |  |
| Age (years) | 48.2<br>(18.0) | 24.1<br>(3.5) | 33.9<br>(3.1) | 44.7<br>(2.8) | 56.8<br>(2.2) | 64.0<br>(3.1) | 73.5<br>(2.2) |
| Body mass index (kg/m<sup>2</sup>) | 24.6<br>(4.3) | 24.4<br>(4.5) | 24.8<br>(4.4) | 25.3<br>(3.8) | 23.6<br>(4.1) | 25.2<br>(4.8) | 24.3<br>(4.7) |
| Body mass (kg) | 66.4<br>(12.3) | 65.9<br>(13.6) | 66.4<br>(13.6) | 69.9<br>(11.5) | 65.2<br>(13.3) | 68.5<br>(9.6) | 62.6<br>(12.7) |
| Height (cm) | 164.2<br>(7.0) | 164.1<br>(7.6) | 163.7<br>(7.5) | 166.1<br>(7.2) | 165.9<br>(7.1) | 165.2<br>(6.2) | 160.3<br>(5.3) |
| **Lifestyle characteristics** |  |  |  |  |  |  |  |
| MVPA (min/day) | 36.6<br>(25.6) | 43.5<br>(24.0) | 39.0<br>(24.1) | 25.9<br>(14.4) | 47.0<br>(18.2) | 39.2<br>(34.5) | 22.0<br>(27.6) |
| Protein intake (g/kg/day) | 1.3 (0.4) | 1.4 (0.5) | 1.6 (0.4) | 1.3 (0.4) | 1.4 (0.4) | 1.2 (0.3) | 1.3 (0.4) |
| **Whole-body composition** |  |  |  |  |  |  |  |
| Lean mass (kg) | 41.7<br>(5.4) | 42.6<br>(7.4) | 41.1<br>(4.6) | 44.6<br>(4.8) | 41.7<br>(4.8) | 41.5<br>(3.9) | 38.5<br>(3.9) |
| Fat mass (kg) | 21.8<br>(9.2) | 20.1<br>(8.1) | 22.5<br>(10.4) | 22.3<br>(9.2) | 20.6<br>(10.2) | 24.3<br>(9.3) | 21.6<br>(9.3) |
| **Thigh composition** |  |  |  |  |  |  |  |
| Thigh area (cm<sup>2</sup>) | 181.5<br>(42.0) | 196.3<br>(49.8) | 185.0<br>(38.2) | 189.1<br>(39.1) | 160.1<br>(18.6) | 180.6<br>(39.5) | 170.7<br>(55.8) |
| Thigh muscle area (cm<sup>2</sup>) | 97.3<br>(19.6) | 107.5<br>(20.8) | 105.9<br>(24.1) | 103.1<br>(12.9) | 87.4<br>(12.4) | 93.2<br>(12.7) | 79.4<br>(15.6) |
| Thigh lean muscle area (cm<sup>2</sup>) | 93.0<br>(17.9) | 104.1<br>(19.3) | 99.4<br>(16.1) | 99.8<br>(13.3) | 83.8<br>(12.3) | 88.2<br>(13.1) | 75.1<br>(14.9) |
| Thigh intramuscular fat area (cm<sup>2</sup>) | 3.7 (2.4) | 3.5 (2.9) | 2.7 (1.6) | 3.3 (2.3) | 3.6 (1.5) | 5.0 (3.0) | 4.3 (2.5) |
| Thigh subcutaneous fat area (cm<sup>2</sup>) | 69.9<br>(31.3) | 73.8<br>(34.4) | 68.6<br>(28.1) | 71.1<br>(32.2) | 58.2<br>(10.9) | 72.1<br>(36.4) | 76.3<br>(41.8) |
| **Quadriceps-specific composition** |  |  |  |  |  |  |  |
| Quadriceps cross-sectional area (cm<sup>2</sup>) | 48.2<br>(9.8) | 54.6<br>(11.0) | 50.9<br>(9.1) | 50.9<br>(6.3) | 44.1<br>(6.7) | 45.5<br>(7.4) | 39.0<br>(8.7) |
| Quadriceps intramuscular Fat CSA (cm<sup>2</sup>) | 65.3<br>(54.3) | 60.4<br>(52.2) | 48.4<br>(47.9) | 55.7<br>(35.4) | 100.0<br>(68.3) | 54.5<br>(41.3) | 77.4<br>(69.1) |

Values are mean (SD) unless otherwise stated. MVPA, moderate-to-vigorous physical activity; CSA, cross-sectional area.

### 3.2 Hormonal effects

The effects of menstrual cycle, contraceptive use and hormone replacement therapy are described in the Supporting Information and Data.

### 3.3 Torque-velocity and power-velocity relationships

All primary torque-velocity and power-velocity outcomes decreased linearly with age (Figure 3A-D). When models were adjusted for quadriceps lean CSA, the yearly age-related decline was partly attenuated for P_MAX_ and T_0_, whereas no notable effects were observed for V_0_ or V_opt_ (Figure 3E).

**Figure 2.**
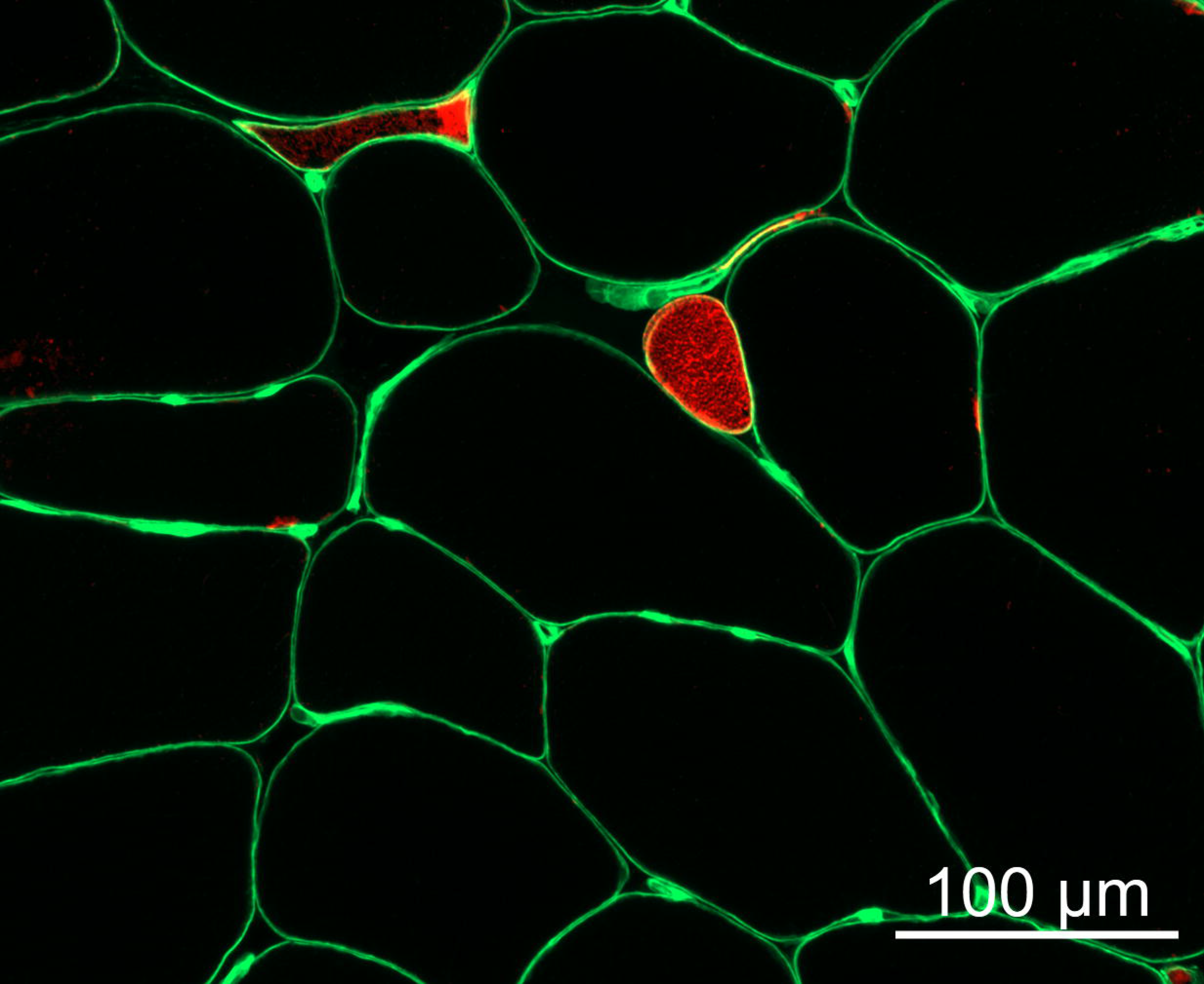
NCAM+ muscle fibers in aged skeletal muscle. Representative image of neural cell adhesion molecule (NCAM, red) and laminin (green). Scale bar=100µm

**Figure 3:**
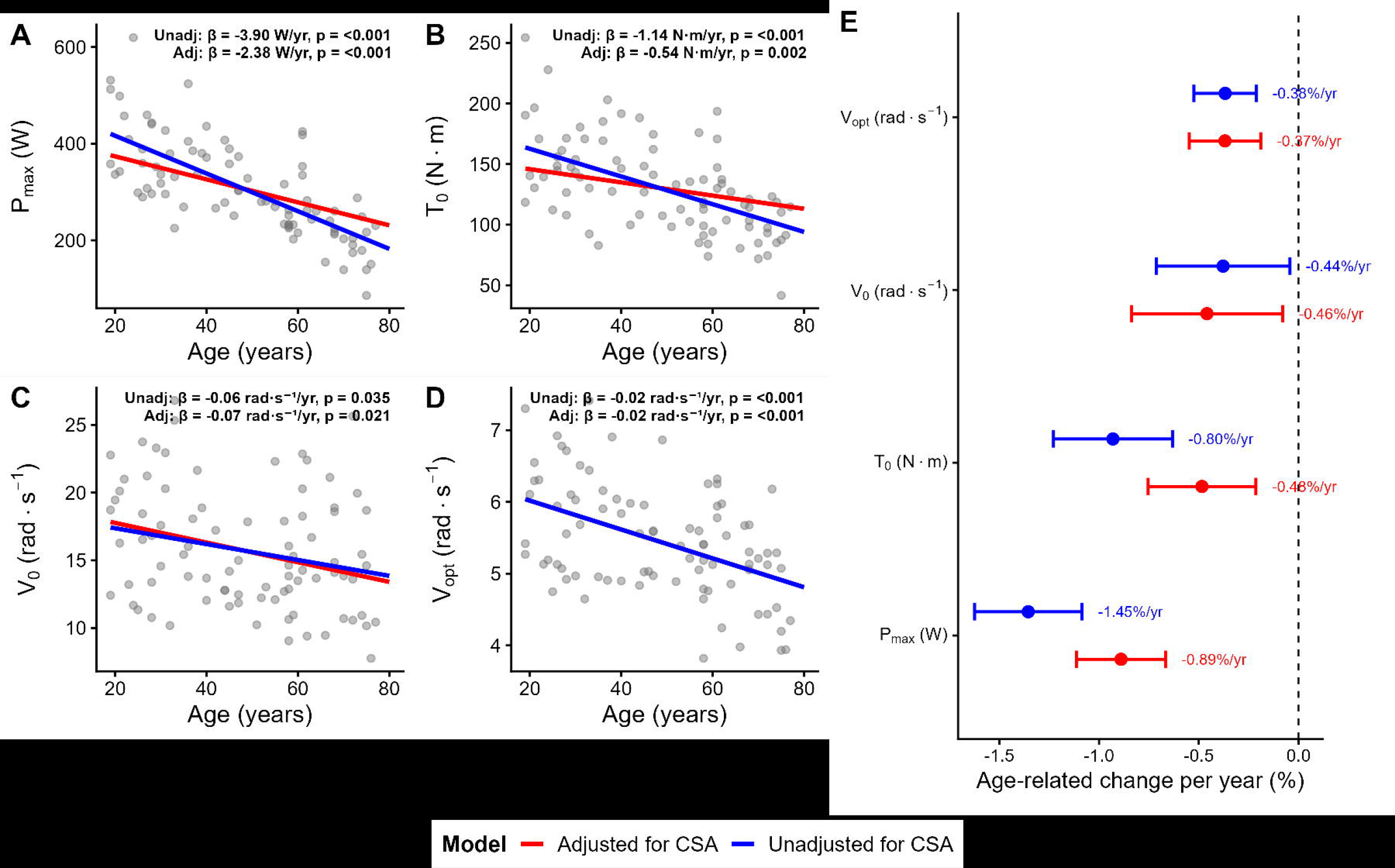
Linear regression models depicting age-related associations for peak power (A), maximal torque (B), maximal velocity (C), and optimal velocity (D) with (red) and without (blue) adjustment for quadriceps cross-sectional area. Panel (E) presents the attenuation of age-related change (% per year) with (red) and without (blue) adjustment for quadriceps cross-sectional area.

### 3.4 Rate of torque development

The relationship between RTD and P_MAX_ are described in the Supporting Information.

Maximum RTD occurred 90.59 ± 2.4ms following torque onset and was not associated with age (R^2^ = 0.012, p = 0.31). RTD during time bins ranging from 0-100ms were isometric, with minor changes in knee angle not associated with age between 100-150ms (0.02 ± 0.02 rad, R^2^ < 0.01, p = 0.7) and 150-200ms (0.07 ± 0.04 rad, R^2^ < 0.001, p = 0.9).

Absolute maximal voluntary RTD decreased non-linearly from 35 years of age (p < 0.0001) and from 43 years of age when adjusted for quadriceps CSA adjusted models (p = 0.003) (Figure 4A). Maximum relative RTD showed no significant age effects either adjusted or unadjusted for quadriceps CSA (p > 0.05) (Figure 4B). A significant age × time bin interaction was observed for absolute RTD (p < 0.0001), indicating that the association between age and RTD differed across time bins. Age-related declines in absolute RTD were observed across all time bins, with slopes of −1.0 N·m·s□¹·year□¹ (95% CI: −2.0, 0.04) at 0–50ms, −4.1 N·m·s□¹·year□¹ (−6.1, −2.1) at 50–100ms, −4.8 N·m·s□¹·year□¹ (−6.0, −3.5) at 100–150ms, and −2.3 N·m·s ¹·year ¹ (−3.4, −1.2) at 150–200ms (Figure 4C). Pairwise comparisons confirmed that the age-related decline was significantly steeper at 50–100ms and 100–150ms compared with 0–50ms, with slope differences of 3.1 and 3.8 N·m·s□¹·year□¹, respectively (both p < 0.001, Figure 3E). The interaction between age and time bin remained significant for relative RTD (p = 0.04), increasing with age during 0-50ms (1.36%·year□¹, 95% CI: 0.1, 2.6) and decreasing with age at 100–150ms (−2.25%·year□¹, 95% CI: −3.7, −0.8) (Figure 4D). Pairwise comparisons indicated that the age-related change was significantly steeper at 100-150ms than at 0–50ms, with a slope difference of 3.6%·year□¹ (p < 0.03). No other differences with age or between time bins were observed (Figure 4F).

**Figure 4.**
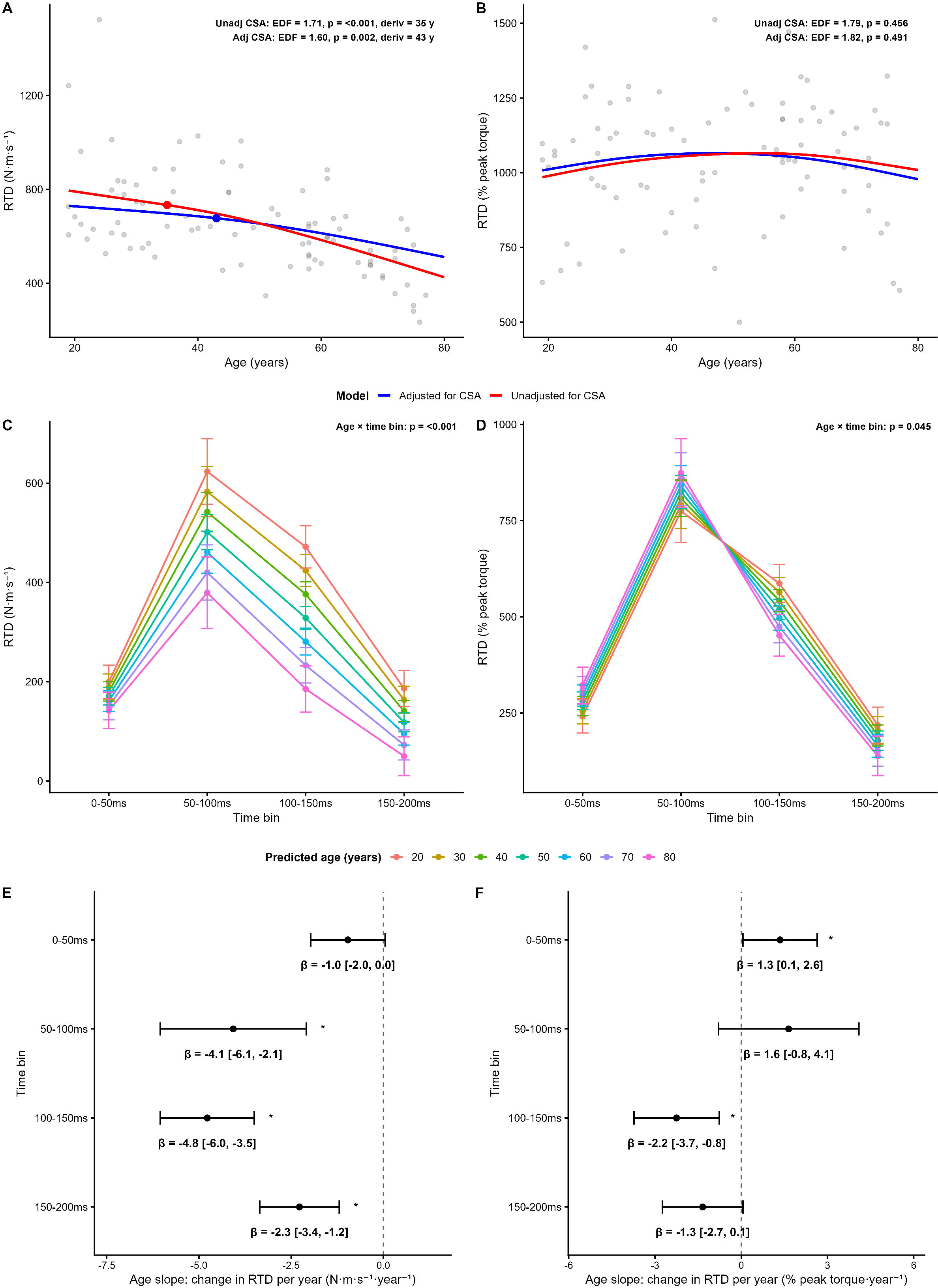
Voluntary absolute RTD (A) and RTD normalised to peak force (B) were modelled using generalised additive models, with (blue line) and without (red line) adjustment for quadriceps cross-sectional area (CSA). Panels C–F show the age × time-bin interaction from mixed models. Panels C and D show estimated marginal means across time bins for absolute RTD and normalised RTD, respectively, across age groups. Panels E and F show the corresponding age-related slopes representing the estimated change in RTD per year of age within each time bin. Error bars represent 95% confidence intervals.

Maximal evoked RTD decreased non-linearly from ∼49 years of age (p = 0.001), however this was attenuated after adjusting for CSA (p = 0.07). No age-related differences were observed for evoked RTD normalised to peak force (p = 0.12), although significant linear reductions were shown following CSA adjustment (p = 0.001).

### 3.5 Electromyography

Generalized linear models revealed significant non-linear trajectories across all time bins for rectus femoris muscle (Figure 5A-C), with comparable or increased average and maximum EMG amplitude during early and mid-adulthood before declining from ∼60 years of age (all p < 0.05), whereas the global average demonstrated a similar trajectory but was not significant (p = 0.08). VL and VM demonstrated a non-linear increase with age during the 0–50ms phase only (VL p = 0.02; VM p = 0.03) whereas no significant changes were observed for later phases or global averages.

**Figure 5.**
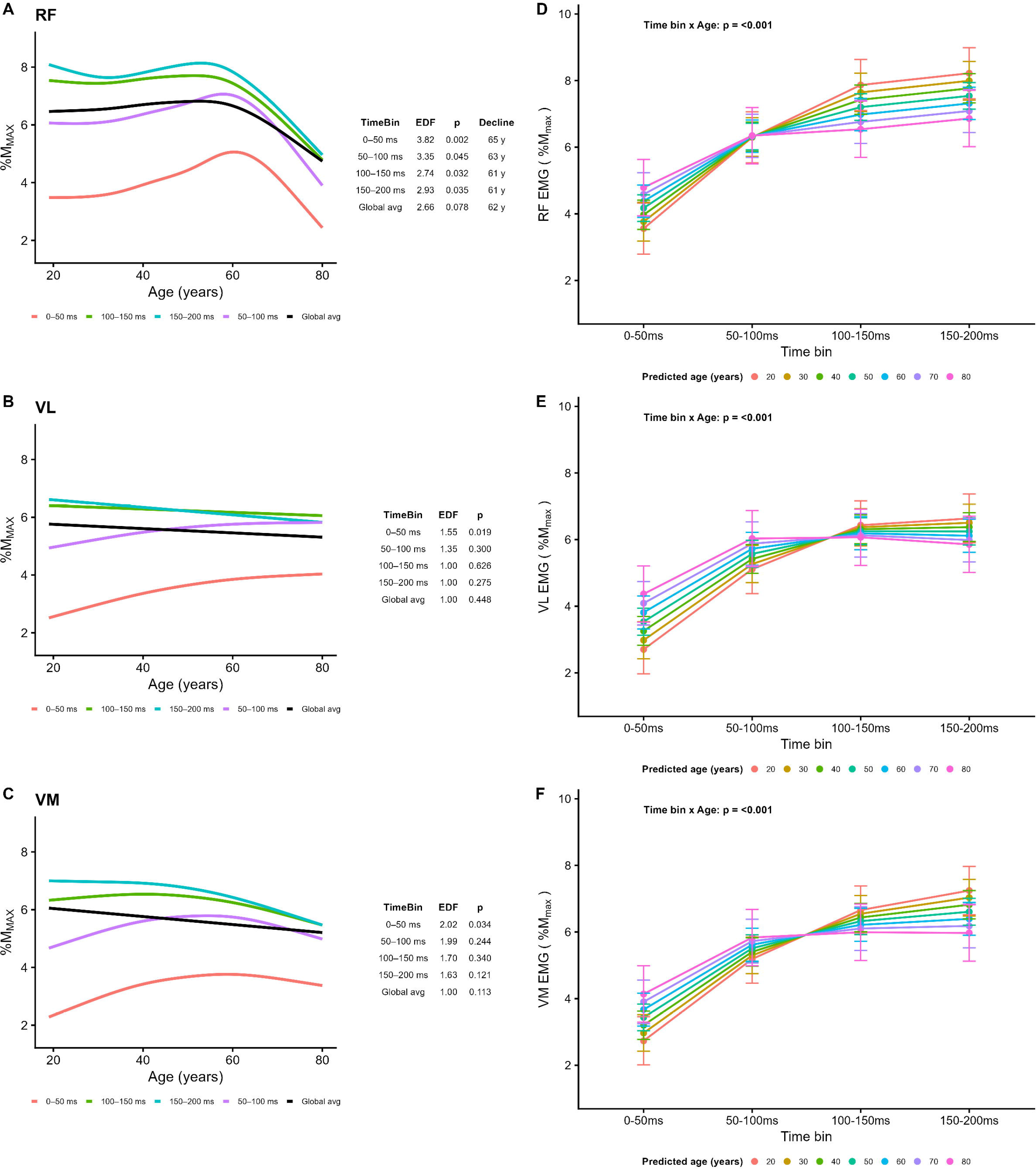
Surface electromyography (EMG) amplitude for rectus femoris (RF), vastus lateralis (VL), and vastus medialis (VM). Panels A–C show model-derived trajectories from generalised additive models illustrating age-related trajectories in average EMG amplitude across time bins (0–50ms, 50–100ms, 100–150ms, and 150– 200ms), with coloured lines representing model-predicted values across the adult lifespan (20–80 years). Panels D–F present estimated marginal means (±95% CI) from mixed models across time bins for each muscle, stratified by age.

Across all quadriceps (RF, VL, VM) significant age x time bin interactions were observed for both average and maximum EMG amplitude similarly (all p < 0.001). The interaction effects for average EMG amplitude are illustrated in Figure 5D-F, where older adults generally displayed higher quadriceps EMG amplitude during earlier time bins < 100ms and lower quadriceps EMG amplitude during later time bins >100ms and <200ms. Rate of EMG rise (RER) decreased linearly with age during 50-100ms for all quadriceps muscles (RF: -0.62 ± 0.13%, p < 0.0001; VL: -0.38 ± 0.12%, p = 0.001; VM: -0.41 ± 0.09%, p < 0.0002), but no changes were shown for 0-50ms time bins (all p > 0.05; Figure 6).

**Figure 6.**
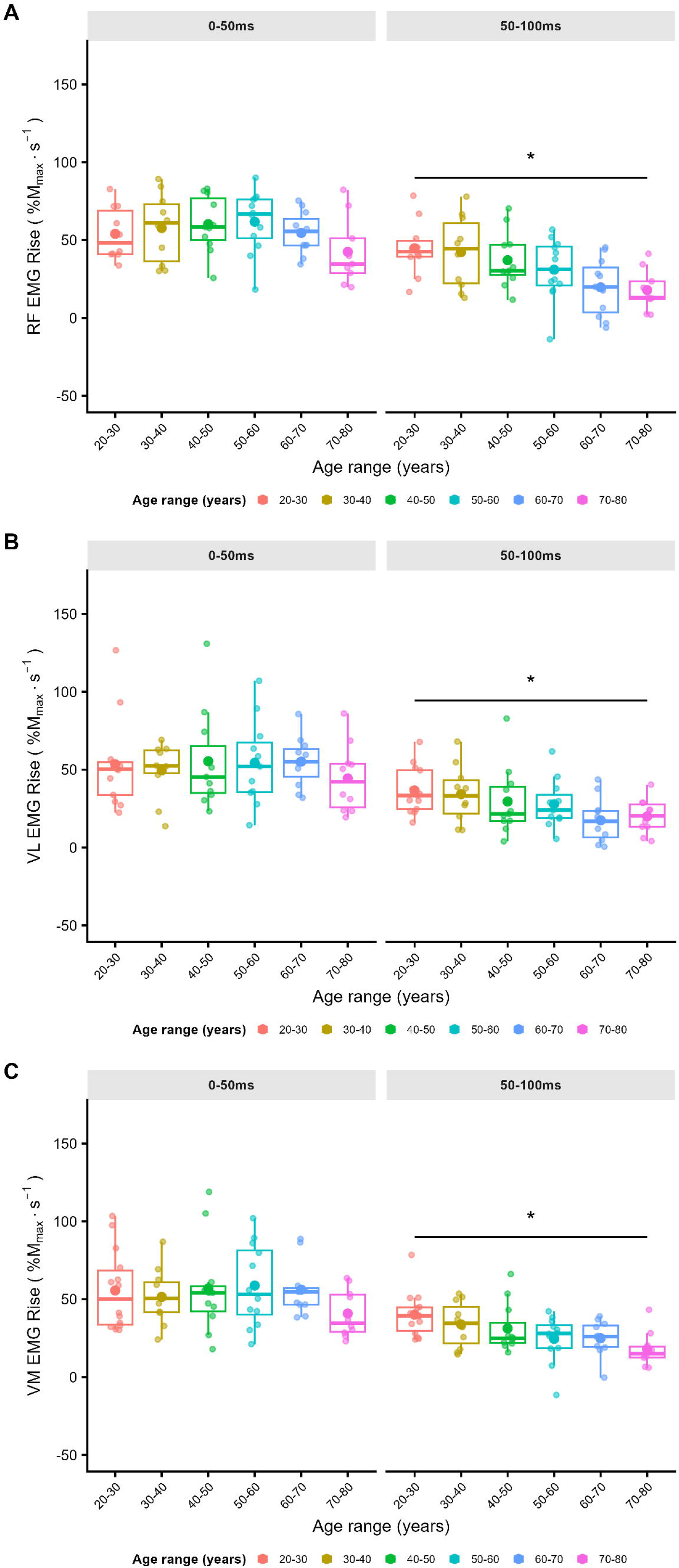
EMG rate of rise across 0-50ms and 50-100ms time bins for RF (A), VL (B), and VM (C). Data is expressed as estimated marginal means (±95% CI) from mixed models across time bins and stratified by age. *Significant age effect (p < 0.05).

### 3.6 Molecular markers of neuromuscular function

For every one-year increase in age, there was a 6.3% increase in the odds of NCAM fibre presence (OR = 1.1, 95% CI: 1.0–1.1, p = 0.01). It was associated with reduced P_MAX_ (β = -0.27 Watts, p = 0.01), T_0_ (β = -0.40 N·m, p = 0.007), V_0_; (β = -0.44 rad.s^-1^, p = 0.02) and V_opt_ (β = -0.37 rad.s^-1^, p = 0.02), although none of these associations remained significant after multiple testing correction (all p = 0.08).

Attenuation analysis revealed that age-related decline in P_MAX_ was modestly attenuated after adjustment for the presence of NCAM (∼14%), and that age-related decline in T_0_, V_0_ and V_opt_ were substantially attenuated after adjustment for the presence of NCAM (∼33%, ∼60% and ∼28%, respectively), where the presence of NCAM independently associated with lower P_MAX_, T_0_, V_0_ and V_opt_.

Following p-value adjustment, age was significantly associated with linear declines in acetylcholinesterase (ACHE) and the β1 subunit of the nicotinic acetylcholine receptor (CHRNB1), and with linear increases in laminin alpha-2 (LAMA2), muscle-specific kinase (MUSK) and the α1 subunit of the nicotinic acetylcholine receptor (CHRNA1) (all FDR<0.05). Of the calcium-handling genes investigated, calsequestrin 1 (CASQ1) and ORAI1, which encodes the pore-forming subunit of the store-operated calcium entry channel, significantly decreased with age (FDR<0.05) (Figure 7). The complete list of neuromuscular and calcium-handling genes investigated and associated outcomes can be found in the Supporting Data.

**Figure 7.**
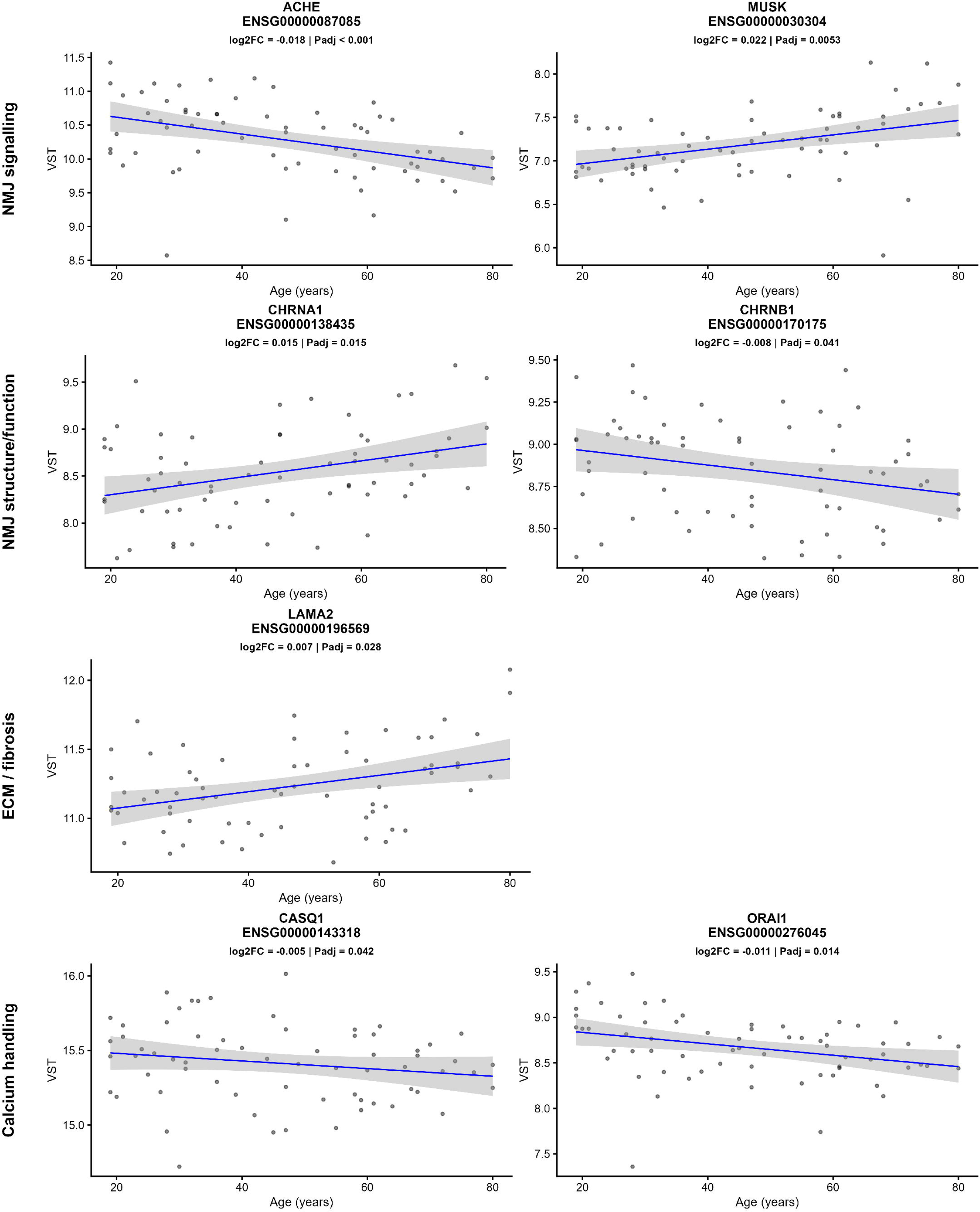
VST-normalised gene expression across age. Points represent individual samples and lines show LM fits (±95% CI). Each panel shows a gene (gene symbol and Ensembl ID), grouped by functional category (NMJ signalling, NMJ structure/function, ECM, and calcium handling). VST: variance stabilised gene expression.

## 4. DISCUSSION

### 4.1 Summary

This study is the first to map the trajectory of age-related changes in quadriceps maximal power across each decade of the female lifespan and to provide mechanistic insight into the role played by skeletal muscle mass, neural drive, denervation and the neuromuscular junction and calcium handling transcriptome. The main findings are: (i) all torque-velocity and power-velocity outcomes decreased linearly with age, with quadriceps lean CSA only partly offsetting the decline in P_MAX_ through mitigating the decrease in T_0_ but not V_0_; (ii) *rectus femoris* was the only quadriceps muscle to exhibit an age-related decline in EMG amplitude, which occurred from ∼60 years of age; (iii) time-bin specific analysis revealed phase-dependent effects whereby, compared to younger females, older females generated more relative RTD and had higher quadriceps neural drive during early time-bins (0-100ms), but lower relative and absolute RTD and neural drive during later time bins (100-200ms); and (iv) NCAM^+^ presence was associated with a global impairment of the force–velocity profile and significantly explained part of the decline in these outcomes beyond the effects of ageing alone.

### 4.2 Torque-velocity and power-velocity relationship

The yearly decline in absolute knee extension power of -3.9 ± 3.8 W was comparable to others who compared power generated during isokinetic contractions between younger and older females [10]. However, after adjusting for quadriceps lean CSA our data showed an attenuation in maximal power by ∼40%, which is much less than previously reported for maximal power normalised to whole-thigh mass (∼85%) [10]. Thus, our data show that non-specific methods may overestimate the contribution of muscle mass to power loss in ageing females and supports a role for other confounding mechanisms. For example, *rectus femoris* was the only superficial quadriceps muscle to display age-related decreases in EMG amplitude which occurred from ∼60 years of age and which were significant during time bins between 100-200ms following force onset and tended to decrease globally. Where we previously suggested that *rectus femoris* may be more vulnerable to denervation across the lifespan compared to the *vasti* [9], here we demonstrate reduced capacity to achieve full volitional recruitment of *rectus femoris* motor units beyond 60 years of age, which may contribute to maximal power loss in older females. *Rectus femoris* is susceptible to reduced activation during faster gait speeds in older compared to younger adults [31] and may partly explain why sarcopenia diagnoses are higher in women when using the International Working Group on Sarcopenia criteria [1]. However, bi-polar surface electromyography has its limitations [32] and more direct investigation into female specific age-related changes in quadriceps motor unit discharge characteristics using high-density surface (HDEMG) and intramuscular (iEMG) techniques is therefore needed.

Maximal torque decreased with age at a greater rate compared to velocity, although after adjustment for quadriceps lean CSA the yearly decline in torque and velocity was similar (Figure 3E). Whereas a decrease in maximal torque is more consistently observed in aging females, the decline in maximal velocity is less clear [6, 10] likely due to differences in assessment methodology (e.g., T-V modelling procedure, isotonic vs. isokinetic vs. unloaded contractions, age treated as continuum vs. as group). *Vastus lateralis* fascicle length may be shorter in older compared to younger females [33] and adjusting for fascicle length can attenuate velocity loss in older males [34], although this remains less clear in ageing females and warrants more investigation [8]. Further, type 2 skeletal muscle fibres with fast contraction velocities were unaffected by age in the same cohort (reported in [20]) although increased fibrosis and hybrid fibre proportion may have contributed to slow maximal shortening velocity and subsequent power output [35].

### 4.3 Rapid torque production

The lack of change in maximal relative rate of torque development previously reported for males [17] and shown here for females, may be due to early compensatory increases in neural drive. Relative rate of torque development was higher during earlier time bins 0-100ms for older females and accompanied by higher EMG amplitude for all quadriceps muscles during 0-50ms and for *vasti* during 50-100ms, whereas relative rate of torque development was lower during later time bins 100-200ms for older females and accompanied by decreased EMG amplitude and rate of EMG rise for all quadriceps muscles. Previous studies in males have analysed RTD during overlapping time bins meaning that these distinct changes may have gone undetected [17], or that discordant effects of ageing on rate of torque development during distinct time bins are evident in females compared to males. Although the reasons for these changes require greater insight, our results may suggest increased initial supraspinal drive followed by reduced motor unit recruitment and discharge rate. The initial increase in EMG amplitude may reflect neural compensation for reduced skeletal muscle quality and/or increased musculotendinous stiffness in older females [9, 33] partly supported here by increased quadriceps fibrosis, intramuscular fat, and reduced relative evoked RTD adjusted for skeletal muscle CSA. The later decrease in EMG amplitude and rate of EMG rise supports a neural impairment in RTD [36] and which may reflect reduced capacity to sustain initial motoneuron firing in response to reduced intrinsic motoneuron excitability [37] and/or neuromuscular junction transmission failure/instability [23, 38]. This interpretation is supported by our statistical models which were adjusted for skeletal muscle mass, moderate to vigorous physical activity level and protein intake, and quadriceps voluntary activation assessed by both femoral nerve and cortical stimulation is unchanged in 18-80 year old healthy females [9, 10]. Further, maximal RTD occurs prior to afferent proprioceptive feedback [14] and dynamic knee extension force does not seem to be significantly influenced by antagonist co-activation in ageing females [33].

### 4.4 NCAM and neuromuscular transcriptome

NCAM (Neural Cell Adhesion Molecule, CD56) is a membrane protein mediating cell-cell adhesion between motor neuron and muscle fibres. The presence of NCAM□ fibres indicate increased axonal guidance and reinnervation, reflecting impaired neuromuscular integrity. NCAM fibres therefore represent a robust marker of denervation in healthy ageing muscle [39, 40]. Here we report that NCAM presence is associated with a global impairment of the whole force–velocity profile (including power, torque and velocity) pointing towards age-related impairment of the entire motor unit. Although these associations did not remain significant following multiple testing correction, the uniform direction and magnitude of effects across outcomes support a role for denervation-reinnervation processes in driving age-associated neuromuscular dysfunction. Further, attenuation analyses reveal that NCAM presence captures a substantial component of the variability typically attributed to “ageing.” In particular, the marked attenuation observed for maximal shortening velocity suggests that age-related reductions in contractile speed may be largely mediated by neuromuscular factors rather than intrinsic muscle properties. These results were supported by a robust molecular signature of NMJ instability. Reduced expression of NMJ maintenance genes (ACHE, CHRNB1) alongside increased expression of genes involved in synaptic remodelling and receptor clustering (MUSK, LAMA2, CHRNA1) was consistent with structural remodelling and compensatory reinnervation of the NMJ in response to progressive neuromuscular instability. The selective downregulation of key calcium-handling genes CASQ and ORAI1 with ageing may reflect impaired sarcoplasmic reticulum Ca² storage and Ca² store replenishment during muscle contraction, further contributing to NMJ instability, although the Ca² handling machinery, including the ATP-dependent calcium pump SERCA (see Supporting Data), otherwise appear mostly preserved.

### 4.5 Hormonal influence

We first confirmed that menstrual cycle phase or OC use did not significantly influence any neuromuscular outcomes (see Supporting Information and Data). Our pilot investigation of the influence of hormone replacement therapy in post-menopausal females (n = 5) revealed that average and maximal EMG amplitudes of both *vasti*, and to a lesser extent the *rectus femoris*, were numerically higher in HRT users across multiple time bins. These differences did not persist after multiple testing correction, but the consistency of the direction of effects across related measures may indicate a potential neurophysiological effect. However, the current study is not powered to robustly assess the impact of HRT, and these findings should not be overinterpreted.

Although we quantified sex hormone levels for all participants in this study (reported in [20]), we did not investigate their association with the reported outcomes, as age-related declines in T-V and P-V outcomes were linear (discordant to isometric forces which accelerated during the menopausal transition [9]). However, the decline in maximal RTD both unadjusted and adjusted for quadriceps CSA decreased non-linearly between 35-43 years of age, and more investigation is required to evaluate how hormonal changes during the menopausal transition may impact motor unit discharge rate as a key determinant of rapid force capacity.

## 5. CONCLUSION

We demonstrate that several mechanisms beyond skeletal muscle mass substantially contribute to age-related declines in dynamic torque-velocity, power-velocity and rapid force capacity across the female lifespan. This primarily resembles a neural degenerative profile evidenced by changes in quadriceps voluntary drive, denervation and neuromuscular junction instability. However, none of the participants fell below defined cut-off criteria for sarcopenia diagnoses including isometric knee extension strength <12.5kg and appendicular skeletal muscle mass < 5.5 kg/m² [2] and more research implementing similar methods is warranted to better understand the clinical relevance of our findings in a sarcopenia context. Future research aimed at evaluating the contribution of female sex hormones to motor unit and neuromuscular junction degeneration and instability, in addition to potential protective effects of various exercise, lifestyle, hormone replacement and pharmaceutical interventions, could mitigate the functional burden facing ageing females.

## ACKNOWLEDGMENTS

The authors would like to thank the participants who volunteered their time and Ms Briana Gatto for her role in managing participant recruitment and coordination during the early phases of the study.

