## Supporting information for "Muscle mass and denervation explain variability in maximal power and rapid force across the adult female lifespan"

**Supplementary Methods**

**Experimental Protocol and procedures**

Visit 1 included assessment of quadriceps maximal torque, velocity, power and rate of torque development with concurrent bi-polar surface electromyography recordings from the superficial quadriceps muscles. Participants sat upright on an isokinetic dynamometer (Universal Pro Single Chair model 850-230, Biodex Medical Systems, United States) with straps securing the thorax and pelvis and hip angle set at 85° flexion. Following a standardized warm-up participants completed maximal dynamic knee extensions against five different isotonic loads (0% MVC, 15% MVC, 30% MVC standard and then adjusted between 45-60% MVC dependent on individual strength capacity) and one isokinetic speed (1.047 rad.s^-1^). Two efforts were performed for each load/speed (⁓3 s apart) and 3 min of rest separated different loads/speed. Participants were instructed to avoid any pre-tension before performing each contraction (visually verified) and to extend their knee as “hard and fast as possible” from starting position of ~110° knee flexion to an approximate straight leg position (~0° flexion). Immediately following each pair of contractions, knee angle was passively moved to 110°, 70° and 30° knee flexion and a resting supramaximal 1 Hz electrical stimulus was applied to the femoral nerve to evoke a maximal muscle compound action potential (M_MAX_) in the superficial quadriceps.

All torque (N.m), angle (rad), velocity (rad.s^-1^) and EMG (mV) data were synchronously acquired at 4000Hz and digitally converted with a Powerlab 8/30 (ADinstruments, Australia) in conjunction with a PC running Labchart software (version 8.1.24, ADinstruments, Australia). All analysis was completed offline using Spike2 software (version 7.13, Cambridge Electronic Design, Cambridge, UK). Concentric torque was gravity offset by multiplying the passive torque generated by the weight of the leg in a straightened position (~0° flexion) by the cosine of the recorded knee angle. An additional power channel (W) was calculated by multiplying torque (N·m) by velocity (rad.s^-1^). For all dynamic knee extension contractions, peak and average torque, power and velocity were extracted between ~110° and 30° flexion (0° = full extension). This offset angle was chosen to avoid increased resistance generated by the braking torque (i.e. cushioning) of the dynamometer toward full knee extension during high-velocity contractions. Surface electromyography signals from *vastus lateralis* (VL), *vastus medialis* (VM) and *rectus femoris* (RF) were filtered using a second order Bandpass Butterworth filter (6 – 500 Hz) (Tillin et al., 2010).

Visit 2 included measures of body composition and blood and muscle sampling. Participants were provided with a standardised meal to consume *ad libitum* the night prior to their visit, and they were instructed to fast from midnight. Whole body composition was evaluated via dual energy X-ray absorptiometry (DEXA; GE Lunar,Madison, WI, USA) and quadriceps tissue composition was evaluated via peripheral quantitative tomography (qPCT; Stratec Medizintechnik GmBH, Pforzheim, Germany). A muscle biopsy was then taken from the *vastus lateralis* using the percutaneous needle technique modified to include suction (Evans *et al.*, 1982). First, the skin was sterilised and anaesthetised with 1% lidocaine. Following this, an incision was made through the skin and fascia. The muscle sample was removed, immediately snap frozen in liquid isopentane, and stored at -80°C until needed for analysis.

**Statistics**

Missing covariate data (MVPA (13.5%), quadricep CSA (16.7%) and protein intake (14.6%)) were imputed using the kNN function in the VIM package [34]. Missingness was not significantly associated with observed variables, supporting a missing completely at random assumption, and imputation quality was assessed using density and strip plots.

Model fit between linear and nonlinear models were compared using Akaike Information Criterion (AIC). Non-linearity and model fit accuracy examined the effective degrees of freedom (EDF) were values near 1 indicated approximately linear relationships and k-index diagnostic to ensure adequate basis dimensions. For significant non-linear models we the first and second derivatives were used to identify regions of increase or decline and turning points. Edge regions comprising 5% of the age range were excluded to reduce boundary instability.

Genes with a mean raw count <10 across samples were excluded prior to downstream analyses. Count data were normalised using DESeq2’s internal size factor estimation method. Variance stabilising transformation (VST) was applied for principal component analysis (PCA) and data visualisation. PCA was used to assess overall sample clustering and to confirm that sequencing batch effects were adequately accounted for following batch adjustment.

**Supplementary Results**

**Rate of torque development**

Maximum RTD occurred 90.59 ± 2.4 ms following torque onset and was not associated with age (R^2^ = 0.012, P = 0.31). RTD during time bins ranging from 0-100ms were isometric, with minor changes in knee angle not associated with age between 100-150ms (0.02 ± 0.02 rad, R^2^ < 0.01, P = 0.7) and 150-200ms (0.07 ± 0.04 rad, R^2^ < 0.001, P = 0.9).

Maximal voluntary rate of torque development was strongly associated with P_MAX_ in unadjusted analyses (RTD_MAX_, R² = 0.63, P < 0.0001). P_MAX_ was also positively associated with RTD across individual time intervals (RTD 0–50ms: R² = 0.30, P < 0.0001, RTD 50–100ms: R² = 0.48, P < 0.0001, RTD 100–150ms: R² = 0.39, P < 0.0001 and RTD 150–200ms: R² = 0.19, P < 0.0001). After adjustment for age, maximal voluntary rate of torque development was positively associated with P_MAX_ (RTD_MAX_: partial R^2^ =0.21, P <0.0001; RTD 0-50ms: partial R^2^ = 0.16, P <0.0001; RTD 50-100ms: partial R^2^ = 0.19, P <0.0001; RTD100-150ms: partial R^2^ = 0.005, P <0.005) but not RTD 150-200ms (partial R^2^ = 0.02, P = 0.09).

**Hormonal effects**

Menstrual cycle phase did not significantly influence any of the outcomes (Supplementary data file, sheet 2: LM_MenstrualPhase). Compared with non-users, intrauterine device (IUD) and oral contraceptive pill (OCP) use were nominally associated with higher normalised RTD between 100-150ms (p = 0.016 and p = 0.048, respectively). IUD use was also nominally associated with lower VL electromechanical delay, while implant use was nominally associated with lower maximum VL EMG amplitude (both p < 0.05). However, none of these associations remained significant following correction for multiple comparisons (all FDR-adjusted p < 0.05; Supplementary data file, sheet 3: LM_Contraceptive method).The use of hormone replacement therapy (HRT; or menopausal hormone therapy, MHT) in post-menopausal participants tended to increase EMG amplitude in quadriceps across several time intervals, particularly in *vastus medialis* however, none remained significant after adjustment for multiple comparisons (all FDR-adjusted p < 0.05; Supplementary data file, sheet 4: LM_HRT). These findings should be interpreted cautiously because only five participants reported HRT/MHT use.
